# Enhanced axonal transport in large vertebrates: KIF5A adaptations in giraffes and pythons

**DOI:** 10.1101/2025.05.04.652084

**Authors:** Taketoshi Kambara, Lu Rao, Yosuke Yamagishi, Kazuho Ikeda, Daisuke Taniguchi, Tsuyoshi Imasaki, Hideki Shigematsu, Naoki Sakai, Arne Gennerich, Ryo Nitta, Yasushi Okada

## Abstract

Efficient axonal transport is essential for neuronal function, particularly in species with exceptionally long axons. The Kinesin-1 motor protein KIF5A plays a key role in this process, but whether and how it adapts to the transport demands of large vertebrates remains unclear. Here, we show that KIF5A from giraffes (GcKIF5A) and pythons (PbKIF5A), moves 25% faster than its mouse counterpart (MmKIF5A) on neuronal microtubules *in vitro* and within axons of cultured mouse hippocampal neurons. This enhanced velocity, driven by three unique amino acid substitutions (R114Q, S155A and Y309F), facilitates long-distance transport in these species and represents a case of convergent evolution. Our structural analysis reveals that accelerated ADP release underlies the increased speed of GcKIF5A. Despite exhibiting a reduced force generation, GcKIF5A maintains efficient cargo transport under load. Furthermore, GcKIF5A exerts less drag in mixed-motor environments, an adaptation particularly beneficial for multiple motor-driven long-distance axonal transport. These findings reveal that KIF5A has evolved specific adaptations to facilitate efficient axonal transport in large vertebrates, highlighting the evolutionary plasticity of kinesin motors.

## INTRODUCTION

Neurons are highly polarized cells with complex dendrite networks and elongated axons. Some axons extend extraordinary distances, such as the human sciatic nerve and motor neurons of the spinal cord, which can reach up to one meter in length. Axonal transport is essential for neurite formation and extension, synaptic function, and neuronal survival, ensuring a continuous supply of mRNAs, proteins, synaptic vesicles, and membrane components from the cell body to the axon tip^1^. Disruption in this transport system is linked to neurodevelopmental and neurodegenerative diseases^2–6^.

Kinesin-1, or KIF5, plays a central role in intracellular transport, particularly in the long-distance axonal transport of diverse cargos^7,8^. Mammals express three KIF5 paralogs: the ubiquitously expressed KIF5B, and the neuron-specific KIF5A and KIF5C^9,10^. KIF5A is essential for neuron survival and axon outgrowth, particularly during development, whereas KIF5C is prominently expressed in motor neurons and supports their maintenance^9,11^. KIF5A and KIF5B knockout mice are lethal, whereas KIF5C knockout mice are viable^9,12,13^. Despite their high sequence conservation, KIF5 isoforms have distinct functional roles, as mutations in KIF5A are associated with neurodegenerative disorders such as Hereditary Spastic Paraplegia (HSP), Charcot-Marie-Tooth Type2 (CMT2), and amyotrophic lateral sclerosis (ALS)^14–20^.

Point mutations in the motor domain of KIF5A cause an autosomal dominant form of HSP, hereditary spastic paraplegia-10 (SPG10), which predominantly affects the longest motor tracts in the spinal cord^14,21–23^. Most HSP-associated mutations impair KIF5A’s motor activity, reducing velocity by 25-75% compared to the wild-type (WT) protein^16^. Because these mutant proteins are typically expressed at levels comparable to WT, axonal transport in heterozygous patients is only moderately slowed, yet this partial impairment selectively affects neurons with the longest axons.

This length-dependent vulnerability in HSP suggests a relationship between axonal length and transport velocity requirements. In the human nervous system, where some motor neurons approach one meter in length, modest velocity reductions correlate with selective degeneration of the longest motor tracts. This clinical observation suggests that long-distance axonal transport may face several challenges: the cumulative time required for transport over extreme distances, the probability of motor detachment from microtubules during transport, and the finite lifespan of transported cargoes. These factors could theoretically become more significant constraints as axonal length increases, suggesting that even modest velocity enhancements might confer advantages in species with extraordinarily long axons.

Given that a modest reduction in velocity can compromise human axonal integrity, large vertebrates with substantially longer axons may require kinesins with at least similarly modest velocity enhancements for maintaining neuronal viability.

To test this hypothesis, we examined KIF5A from giraffes, which possess exceptionally long axons. In the tallest giraffes, the neck reaches 2-3 meters, making the total length of their recurrent laryngeal nerves (*N. laryngeus recurrens*) nearly 5 meters^24^ **(Figure 1A)**. The motor tracts of giraffes exceed their neck length, resulting in axons at least three times longer than those in humans and nearly 100 times longer than in mice.

**Figure 1.**
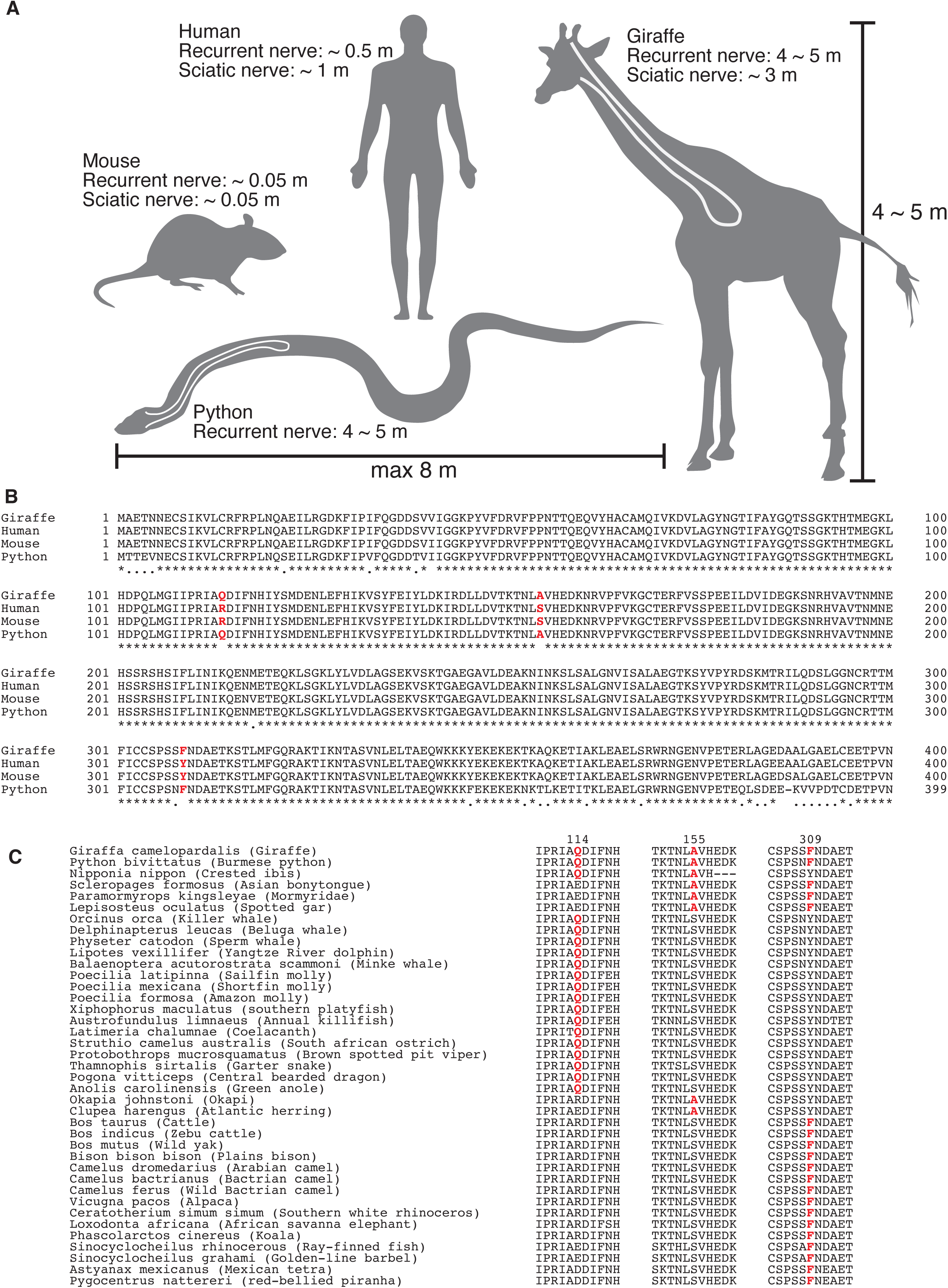
Alignment of KIF5A Sequences Reveals Shared Amino Acid Substitutions in Large Animals with Exceptionally Long Axons. (A) Schematic illustration of axon length in representative species, highlighting the extreme length in giraffes and Burmese pythons. (B) Sequence comparison of the N-terminal motor domain of KIF5A (amino acids 1-360) from human, mouse, giraffe (GcKIF5A), and Burmese python (PbKIF5A). Residues at positions 114, 155, and 309—unique to giraffe and python—are highlighted in red. (C) Multiple sequence alignment of the regions surrounding residues 114, 155, and 309 from 39 selected species (out of 150 analyzed) that differ from both human and mouse at these sites. The scientific and common names are shown for each species. The triple substitution observed in GcKIF5A and PbKIF5A was not present in the closely related okapi— which has a short neck—and was also absent in shorter snakes.

Here, we show that giraffe KIF5A (GcKIF5A) moves ∼25% faster than mouse KIF5A along taxol-stabilized microtubules composed of porcine brain tubulin and within axons of cultured mouse hippocampal neurons, while no significant velocity difference was observed in dendrites and non-neuronal cells. The motor domain of GcKIF5A differs from MmKIF5A and human KIF5A (HsKIF5A) by only three amino acids. Remarkably, the same substitutions were identified in Burmese python KIF5A (PbKIF5A) among 151 species analyzed, and PbKIF5A also exhibited significantly faster movement than MmKIF5A on microtubules *in vitro*. These findings suggest that in species with axons several meters or longer, KIF5A has evolved to support faster long-distance transport.

Our structural analysis suggests that each of these three amino acid substitutions contributes to a distinct adaptation that enhances motor velocity, optimizes performance under load, and facilitates cooperative transport with multiple motors. This combination of modifications enables efficient long-distance axonal transport—a feature particularly advantageous in large vertebrates with exceptionally elongated axons.

## RESULTS

### Common Amino Acid Substitutions in KIF5A of Giraffes and Pythons

We identified the sequence of giraffe GcKIF5A from the whole-genome shotgun contigs of Masai giraffe (*Giraffa camelopardalis*)^25^ using the tblastn program with the amino acid sequence of MmKIF5A (**Figure 1B** and **S1**). Compared to MmKIF5A and human HsKIF5A, GcKIF5A harbors three unique substitutions in its motor domain (R114Q, S155A, and Y309F). Alignment of these positions across species revealed that they are highly conserved (**Figure S2**). Positions 155 and 309 are typically occupied by Serine or Alanine and Tyrosine or Phenylalanine, respectively. In contrast, position 114 exhibits greater variability, with residues including Glutamine, Glutamic acid, Arginine, or Aspartic acid.

While single or paired substitutions occasionally appeared in cetartiodactylas and certain fish species, only Burmese python KIF5A (*Python bivittatus*, PbKIF5A) shares all three substitutions with GcKIF5A among all 151 species analyzed (**Figure 1C**). Burmese pythons can grow over 8 meter in length, with their recurrent laryngeal nerve exceeding 4 meters in largest individuals, as their heart is positioned 15-25% of the body length from the head^26^ (**Figure 1A)**. Notably, this triple substitution was absent in KIF5A of the okapi, the giraffe’s closest relative with a short neck, as well as in shorter snakes (**Figure 1C** and **S2**).

### GcKIF5A and PbKIF5A Move Faster Than MmKIF5A *In Vitro*

To assess the motile properties of GcKIF5A and PbKIF5A, we expressed and purified recombinant K560-EGFP-tagged proteins and compared them to mouse MmKIF5A (**Figure 2A**, **2B** and **2C**). Tetramethyl rhodamine-labeled microtubules, polymerized from porcine brain tubulin and stabilized with Taxol, were immobilized on a coverslip surface. The movement of individual GFP-tagged KIF5A motors was tracked using total internal reflection fluorescence (TIRF) microscopy.

**Figure 2.**
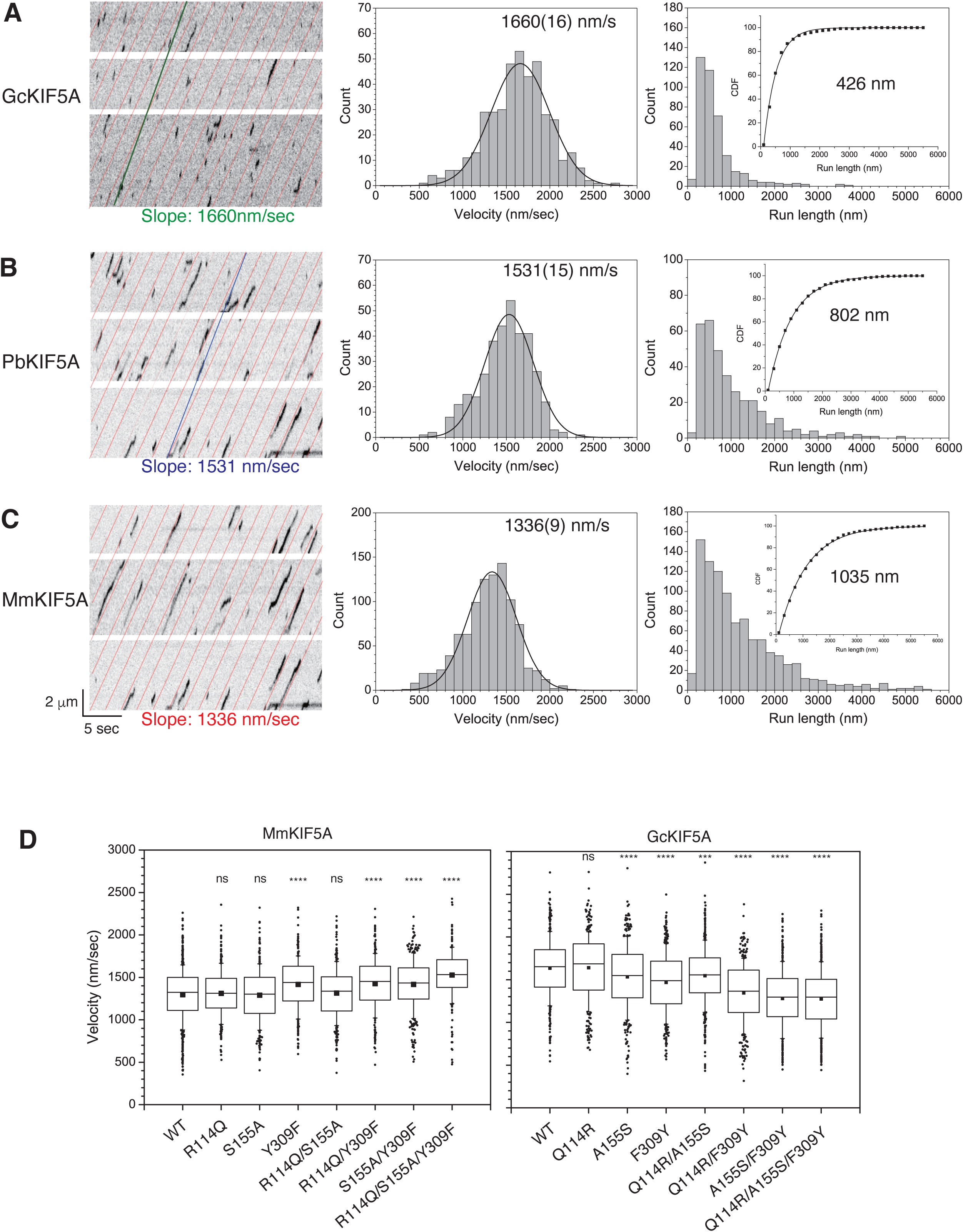
GcKIF5A and PbKIF5A Exhibit Higher Velocities than MmKIF5A on Purified Neuronal Microtubules. GFP-tagged KIF5A motor domain (K560-EGFP) were visualized moving along taxol-stabilized microtubules assembled from porcine brain tubulin. (A-C) Representative kymographs of K560-GFP from giraffe (GcKIF5A), python (PbKIF5A), and mouse (MmKIF5A). The red diagonal lines represent the mean velocity of MmKIF5A (1336 nm/s), blue line shows PbKIF5A (1531 nm/s), and the green line indicates GcKIF5A (1660 nm/s). Mean velocities were quantified from 3-4 independent experiments: MmKIF5A (n=975), PbKIF5A (n=346), and GcKIF5A (n=413). Velocity distributions were fit with a Gaussian function, and values are reported as mean ± SEM. Statistical comparisons were made using Dunnett’s test; P<0.0001 for both PbKIF5A and GcKIF5A versus MmKIF5A. Inset: Average run lengths were determined by fitting cumulative distributions with a single exponential decay function. (D) Effect of GcKIF5A/PbKIF5A-specific amino acid substitutions on motor velocity. K560-EGFP swap mutants containing one or more of the giraffe-specific residues were assayed for motility on taxol-stabilized microtubules. The Y309F alone partially increased the velocity of MmKIF5A. Notably, this residue is also phenylalanine in several other large mammals, including elephant, white rhinoceros, and camel. Data are presented as box-whisker plots: the mean is shown as a filled square, the box represents the 25-75th percentile with a line at the median, and whiskers indicate 10th-90th percentile. Outliers are shown as individual dots. ns, not significant; *** P < 0.001; **** P < 0.0001 versus wild-type KIF5A (Dunnett’s test).

GcKIF5A and PbKIF5A moved significantly faster than MmKIF5A, with velocities of 1660 ± 16 nm/s (mean ± SEM) and 1531 ± 15 nm/s, respectively, representing ∼25% and ∼15% increases relative to MmKIF5A (1336 ± 9 nm/s). Despite their increased velocities, GcKIF5A and PbKIF5A exhibited shorter run lengths than MmKIF5A (**Figure 2A**, **2B** and **2C**). Since KIF5A motors have been reported to display enhanced processivity in viscous solutions containing methylcellulose^27^, we examined motility in the presence of 0.1% Methocel (**Figure S3**). The viscosity under these conditions ranges from 60 – 90 mPa·s, encompassing local cytoplasmic viscosities (80 mPa·s) and cellular cytoplasmic viscosities (1-2 mPa·s)^28–31^. We found that the run length of MmKIF5A tripled as a result of the addition of methylcellulose, whereas GcKIF5A showed only a twofold increase (**Figure S3**). These results suggest that GcKIF5A remains less processive than MmKIF5A even within a cellular environment.

To assess the impact of these three amino acid substitutions shared by giraffe and python KIF5A on its motor activity, we measured the velocities of swap mutants in which the giraffe sequence was introduced into MmKIF5A and vice versa. A single Y309F mutation increased wild-type MmKIF5A velocity from 1336 ± 9 nm/s to 1450 ± 15 nm/s, while the reverse mutation (F309Y) in GcKIF5A decreased its wild-type velocity of 1660 ± 16 nm/s to 1482 ± 18 nm/s. When all three giraffe-specific residues were swapped together (triple mutation R114Q/S155A/Y309F), MmKIF5A velocity was further enhanced to 1545 ± 14 nm/s, while the reverse triple mutation in GcKIF5A further reduced its velocity to 1284 ± 13 nm/s (**Figure 2D)**. These results indicate that all three substitutions collectively contribute to KIF5A’s motor velocity.

In vertebrates, Kinesin-1 consists of two neuron-specific isoforms (KIF5A and KIF5C) and one ubiquitous isoform (KIF5B). To determine whether the increased velocity is specific to KIF5A, we measured the velocities of KIF5B and KIF5C from giraffes and pythons. The sequences of GcKIF5B and GcKIF5C were identified from the whole-genome shotgun contigs of Masai giraffe (*Giraffa camelopardalis*) and compared with their mouse, human, and python counterparts (Supplemental Figs. 4 and 5). Amino acids at positions 113, 154, and 307 in KIF5B, corresponding to positions 114, 155, and 309 in KIF5A, were highly conserved among species. The velocities of KIF5B of mouse, python, and giraffe ranged from 1140 to 1208 nm/s, showing no significant differences among species (**Figure S6**). Similarly, KIF5C exhibited conservation of the three key residues across species. However, all KIF5C isoforms naturally contain F309. Introducing F309 into MmKIF5A increased its velocity (1336 vs. 1450 nm/s), yet MmKIF5C itself showed minimal velocity differences relative to MmKIF5A and exhibited no significant interspecies variation (**Figure S6**). These findings suggest that the increased velocity is a unique adaptation of KIF5A in species with long axons, rather than a general feature of kinesin-1 isoforms.

### GcKIF5A Generates Less Force Than MmKIF5A

KIF5A’s ability to generate force is critical for long-distance cargo transport along axons. To determine whether the three amino acid substitutions in GcKIF5A affect its function under load, we performed optical trapping assays. The duration of motor stalling, or the time KIF5A sustained maximal force generation, was independent of the KIF5A source or motor domain mutation (**Figure S7A)**. However, GcKIF5A exhibited a slightly lower stall force (4.7 ± 0.6 pN) than MmKIF5A (5.3 ± 0.6 pN) (**Figure 3A, 3B** and **S7**). Mutating the three residues in MmKIF5A to match GcKIF5A reduced the stall force to 4.8 ± 0.6 pN (**Figure 3C)**, comparable to WT GcKIF5A. Conversely, introducing the MmKIF5A sequence into GcKIF5A increased the stall force to 5.0 ± 0.5 pN (**Figure 3D)**.

**Figure 3.**
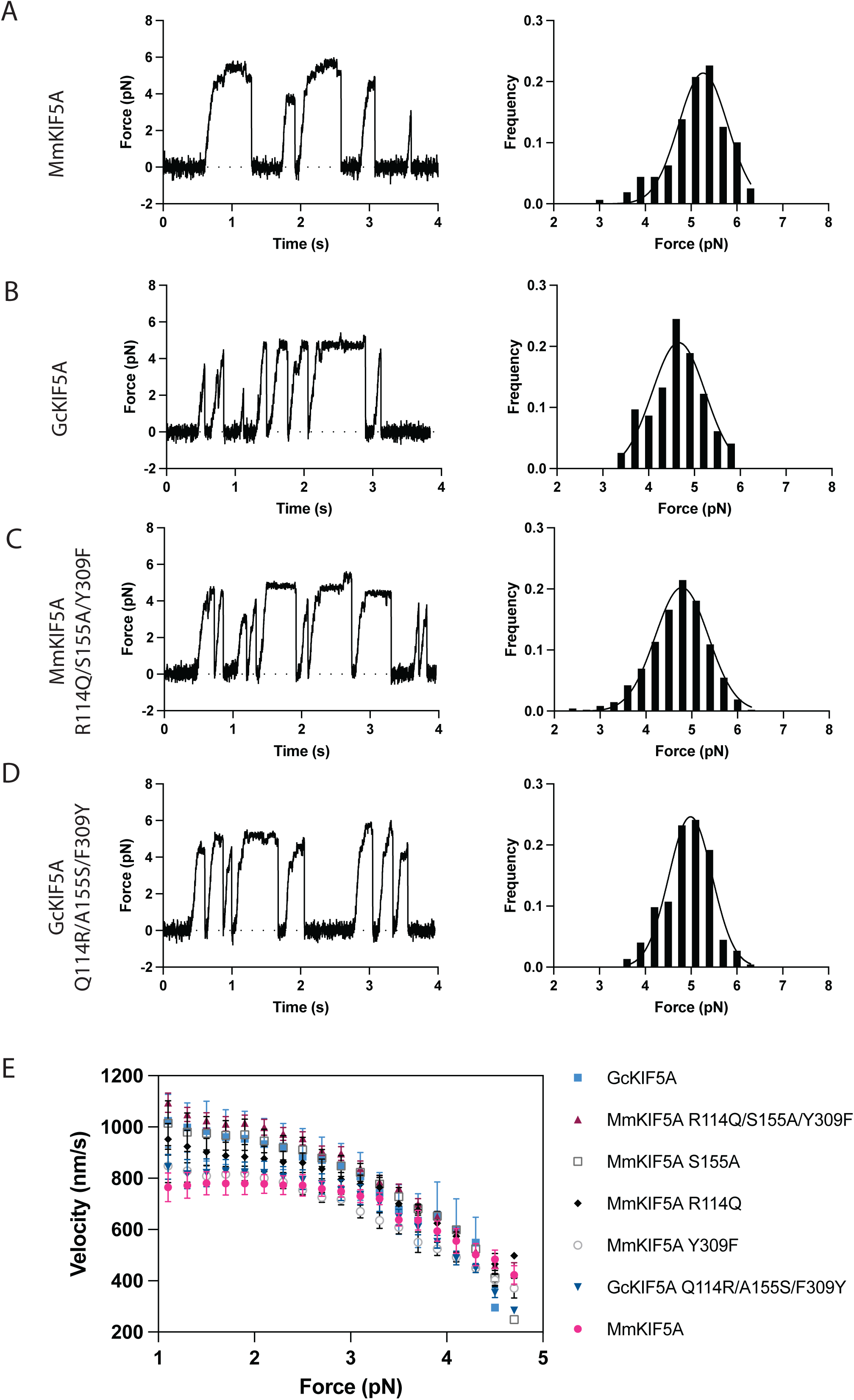
Stall Forces and Force–Velocity Relationships of KIF5A Constructs. **(A-D)** Representative traces and stall force histograms for various KIF5A constructs. Stall force histograms were fitted using Gaussian distributions. **(A)** MmKIF5A, 5.3 ± 0.5 pN (mean ± SD), n = 159; **(B)** MmKIF5A R114Q/S155A/Y309F, 4.8 ± 0.6 pN, n = 476; **(C)** GcKIF5A, 4.7 ± 0.6 pN, n = 196; **(D)** GcKIF5A Q114R/A155S/F309Y, 5.0 ± 0.5 pN, n = 224. **(E)** Force–velocity relationships of the KIF5A constructs. Data were obtained from 4–5 beads per construct; error bars represent the standard error of the mean (SEM).

To assess the effects of individual mutations on force generation, we measured the stall force of MmKIF5A bearing R114Q, S155A, or Y309F substitutions. The resulting forces were 5.0 ± 0.5 pN, 4.8 ± 0.5 pN, and 5.1 ± 0.5 pN, respectively, compared to 5.5 ± 0.5 pN for wild-type MmKIF5A (**Figure S7B)**. While R114Q and S155A had no impact on velocity in the absence of load (**Figure 2D)**, both mutations led to a slight reduction in the motor’s stall force. Additionally, Y309F, which increased the velocity of MmKIF5A (**Figure 2D)**, also weakened force generation. These results suggest that the amino acid substitutions that enhance motor velocity may do so at the expense of reduced force generation, possibly reflecting changes in microtubule binding and/or the power stroke mechanisms.

To understand the functional implications of these adaptations in cellular contexts, we measured the force-velocity relationships for each construct. Based on the Stokes equation, a 100 nm vesicle moving at 1000 nm/s through cytoplasm with viscosity ∼1000-fold higher than water would experience approximately 1 pN of drag force—a physiologically relevant load for axonal transport. While GcKIF5A generated less force than MmKIF5A (4.7 vs. 5.3 pN, **Figure 3A, 3B**), it moved significantly faster under load. For example, at 1 pN load, GcKIF5A moved at over 1,000 nm/s, while MmKIF5A advanced at less than 800 nm/s. Consistent with velocity measurements in the absence of load (**Figure 2D)**, the triple mutant of MmKIF5A also moved at 1,000 nm/s under 1 pN load (**Figure 3E)**. Mechanistically, among the three substitutions, S155A emerged as the dominant contributor to enhanced velocity under load, whereas Y309F, despite its significant effect on unloaded velocity, had a minimal impact under force (**Figure 3E)**. Together, these results suggest that GcKIF5A has evolved to prioritize high-velocity transport under load, effectively powering long-distance axonal transport in large vertebrates.

### Structural Studies Indicate Faster ADP Release from GcKIF5A

To elucidate the molecular basis of GcKIF5A’s enhanced velocity, we employed complementary structural approaches. X-ray crystallography provided high-resolution (1.85-1.95 Å) insight into the nucleotide-binding pocket in the absence of microtubules. In comparison, cryo-electron microscopy (cryo-EM) revealed the motor-microtubule interface at moderate resolution (4.5 Å). This combination allowed us to capture both the intrinsic structural features of GcKIF5A and the conformational changes induced by microtubule binding.

Gln114, Ala155, and Phe309 in GcKIF5A are positioned at key structural sites: helix α2 above the nucleotide-binding pocket, the beta-strand β5a facing the microtubule-binding interface, and the root of helix α6 near the nucleotide-binding pocket, respectively. These residues are expected to influence the mechanochemical cycle of KIF5A. To investigate their contributions, we solved the crystal structures of the GcKIF5A motor domain in the presence of 10 mM ADP or 10 mM AMP-PNP (adenylyl-imidodiphosphate) at 1.95 Å or 1.85 Å resolution, respectively (**Table S1**, **Figure 4A)**. Both structures lacked nucleotides in the binding pocket, instead containing a sulfate ion from the crystallization buffer, which stabilized the P-loop and prevented deformation. While nucleotide-free KIF5 structures have been reported previously^32,33^, EDTA or apyrase is typically required to accelerate the ADP release—the rate-limiting step for KIF5 in the absence of microtubules or tubulin. However, GcKIF5A has no nucleotides in the pocket even in the absence of EDTA or apyrase, suggesting a naturally faster ADP release step.

**Figure 4.**
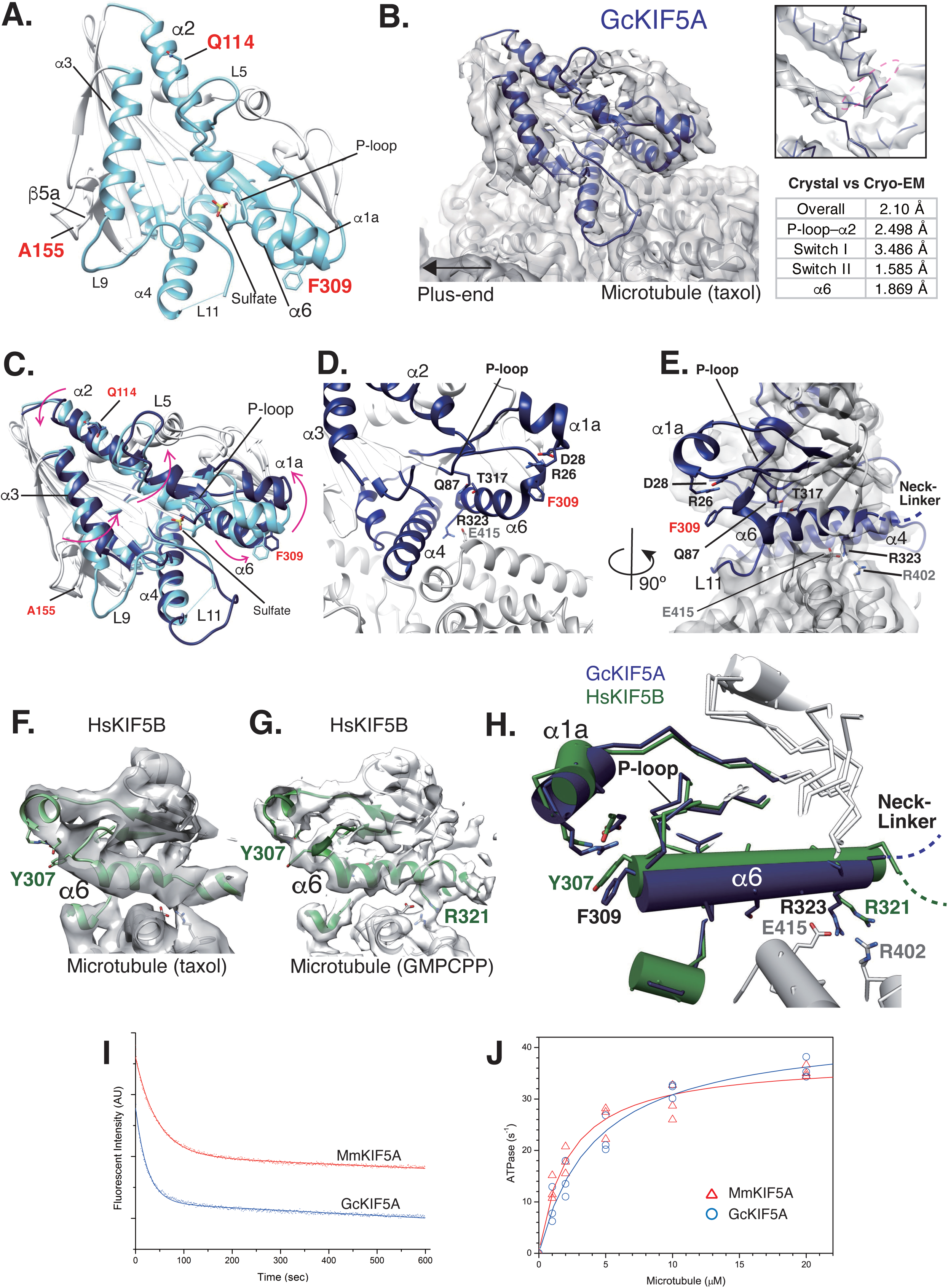
Structural and Biochemical Analysis of the GcKIF5A Motor Domain. (**A)** Crystal structure of GcKIF5A in the nucleotide-free state. (**B)** Cryo-EM structure of GcKIF5A in the nucleotide-free state bound to taxol-stabilized GDP microtubules. The upper right panel shows the empty nucleotide-binding pocket; the lower right lists RMSD values comparing GcKIF5A crystal and cryoEM structures. (**C)** Conformational changes in GcKIF5A upon microtubule binding, highlighting rearrangements in key switch regions. (**D-E)** Detailed views of the cryo-EM structure of GcKIF5A showing interactions among the nucleotide-binding P-loop, helix a1a, and helix a6, and the contact between a6 and a-tubulin. (**F)** Cryo-EM structure of human KIF5B (HsKIF5B) in the nucleotide-free state)bound to taxol-stabilized GDP microtubules (PDBID: 3J8X, EMD-6187). (**G)** Cryo-EM structure of HsKIF5B in the nucleotide-free state bound to GMPCPP microtubules (PDBID: 6OJQ, EMD-20092). (**H)** Structural comparison of GcKIF5A and HsKIF5B in the region surrounding helix a6, revealing conformational differences linked to motor-microtubule interactions. (**I)** Microtubule-stimulated ATPase activity of GcKIF5A and MmKIF5A, showing a higher catalytic rate for GcKIF5A. (**J)** ADP release kinetics from GcKIF5A and MmKIF5A, demonstrating faster nucleotide exchange by GcKIF5A.

The residues in MmKIF5C (PDB ID: 3WRD) and HsKIF5B (PDB ID: 5LT0 and 5LT2) corresponding to GcKIF5A’s unique substitutions are His114, Ala155, and Phe309 in MmKIF5C, and Gln113, Ser154, Tyr307 in HsKIF5B (**Figure S8**). In the absence of microtubules, the H114Q, A155S, and F309Y mutations did not produce substantial conformational changes in GcKIF5A. Kinesin-1 structures have been reported in distinct conformational states: ATP-like (3WRD, MmKIF5C), ADP-like (5LT0, HsKIF5B), and apo-like (5LT2, HsKIF5B)^32,33^ (**Figure S8A)**. Based on root-mean-square deviations (RMSDs), GcKIF5A aligns most closely with the ADP-like conformation (PDB ID: 5LT0, RMSD = 2.0 Å), although switches I and II adopt distinct conformations, likely due to crystal packing or buffer conditions. Notably, the P-loop groove is occupied by a sulfate ion from the crystallization buffer, which stabilizes the P-loop and influences the positioning of helix α2 and loop L5. In Kinesin-5, L5 covers the nucleotide and regulates the ATPase activity; in contrast, L5 in GcKIF5A is flipped away from the nucleotide-binding pocket, potentially facilitating faster ADP release.

We next determined the cryo-EM structure of GcKIF5A bound to taxol-stabilized porcine brain GDP-microtubules at 4.5 Å resolution in the apo state (**Table S2**, **Figure S9, 4B)**. The structure revealed no nucleotide in the nucleotide-binding pocket (inset of **Figure 4B)**. Comparison of the nucleotide-free GcKIF5A structure with and without microtubules (**Figure 4C)** showed the expected conformational rearrangements in switches I and II upon microtubule binding. Specifically, switch I α3 rotated counterclockwise, and switch II L11 became stabilized. Coupling between the P-loop and helix α2 with helices α1a and α6 was supported by hydrophilic interactions: Gln87 (P-loop) with Thr317 (α6), and Phe309 (α6) with Arg26 (α1a) (**Figure 4D, 4E**). Additionally, helix α6 interacted with α-tubulin through Arg323 (α6) and Glu415 (H12), suggesting that microtubule binding is structurally relayed from α6 to the nucleotide-binding pocket.

We next compared the cryo-EM structure of nucleotide-free GcKIF5A bound to taxol-stabilized GDP microtubules with the published structures of nucleotide-free HsKIF5B bound to taxol-stabilized GDP microtubules (**Figure 4F)** or GMPCPP microtubules (**Figure 4G)**. In GcKIF5A, Ala155, located on β5a and facing the microtubule-binding interface, did not appear to contribute directly to microtubule binding. The corresponding density for β5a, including A155, was very weak, indicating conformational flexibility (**Figure S8B)**. In contrast, Ser155 in HsKIF5B also lies away from the microtubule-interface, but the β5a density in HsKIF5B was stronger, suggesting the hydrophilic nature of Ser155 stabilizes this region, potentially contributing to microtubule binding (**Figure S8C)**. The residue corresponding to Gln114 in HsKIF5B was also glutamine, and no significant structural differences were observed at that site. In contrast, the substitution of Phe309 in α6 of GcKIF5A for Tyr307 in mouse and human KIF5A/B markedly altered the microtubule-binding properties. The hydroxy group of Tyr307 in Mm/HsKIF5A/B influenced the coupling between the P-loop, α1a, and α6, whereas Phe309 in GcKIF5A allowed tighter packing between these elements, potentially stabilizing the nucleotide-binding region and modifying the motor’s microtubule interaction (**Figure 4F, 4G** and **4H**).

As described above, helix α6 of GcKIF5A interacts with α-tubulin through a salt bridge between Arg323 (GcKIF5A) and Glu415 (α-tubulin) (**Figure 4D, 4E**). In contrast, the corresponding Arg321 in HsKIF5B is oriented to the right due to α6 movement and is unable to bind to Glu415 (**Figure 4F, 4G** and **4H**). Additionally, Arg402 of α-tubulin is positioned between Arg321 of α6 and Glu415 of α-tubulin, further preventing salt-bridge formation. As a result, α6 in GcKIF5A is positioned closer to the microtubule surface, providing greater structural stabilization. Given that helix α6 forms the base of the neck-linker, its stabilization likely supports more effective neck-linker docking and undocking movements. Moreover, because α6 is directly coupled to the nucleotide-binding pocket, its microtubule-induced stabilization may also facilitate more efficient ADP release from GcKIF5A.

Supporting our structural observations, the rate of ADP release from GcKIF5A (0.047 ± 0.005 s^−1^) was approximately twice that of MmKIF5A (0.023 ± 0.001 s^−1^)^34^ (**Figure 4I)**. In addition, the microtubule-stimulated ATPase activity (*k_cat_*) of GcKIF5A was significantly higher (43.9 ± 2.4 s^−1^, with a K_1/2,MT_ of 4.2 ± 0.7 μM) compared to MmKIF5A (*k_cat_* = 37.7 ± 1.6 s^−1^, with a K_1/2,MT_ = 2.2 ± 0.3 μM) (**Figure 4J)**. These kinetic differences support the conclusion that GcKIF5A’s higher velocities under unloaded and loaded conditions arise from an accelerated catalytic cycle, driven by faster ADP release. To further investigate the physiological relevance of these adaptations, we next examined the behavior of these motors in living neurons.

### GcKIF5A Moves Faster Than MmKIF5A in Axons, but Not in Dendrites and Non-Neuronal Cells

Previous studies have shown that MmKIF5C preferentially moves along microtubules in neuronal axons^35^. Kinesin motility is influenced by the chemical and biophysical properties of microtubules, including tubulin isotype composition, post-translational modifications, nucleotide state, microtubule-associated proteins (MAPs), and stabilization agents such as taxol^36,37^. Because neuronal microtubules are enriched with specific post-translational modifications and neuron-specific isotypes^38^, we hypothesized that GcKIF5A’s faster velocity may depend on the distinct structural features of axonal microtubules. To test this, we compared the velocities of GcKIF5A and MmKIF5A in cultured mouse hippocampal neurons, which provide well-defined axonal and dendritic compartments, as well as in non-neuronal cells as controls.

We transfected mouse hippocampal neurons with a K560-EGFP construct of GcKIF5A or MmKIF5A and measured motor velocities. MmKIF5A moved at similar speeds in both axons and dendrites, whereas GcKIF5A exhibited compartment-specific behavior—moving significantly faster in axons than in dendrites. In axons, GcKIF5A was approximately 20% faster than MmKIF5A, while in dendrites, both MmKIF5A and GcKIF5A moved at comparable velocities (**Figure 5A)**. Treatment with taxol eliminated this compartment-specific advantage of GcKIF5A by increasing its velocity in dendrites to match its axonal speed, while having minimal effect on MmKIF5A velocities in either compartment (**Figure 5A)**. These findings are consistent with our *in vitro* results using taxol-stabilized microtubules assembled from porcine brain tubulin, where GcKIF5A also exhibited higher velocities than MmKIF5A (**Figure 5A)**. In contrast, when we measured motor velocities in mouse cortical astrocytes and mouse embryonic fibroblasts (MEFs), no significant differences were observed between GcKIF5A and MmKIF5A (**Figure 5B)**. Together, these results suggest that the increased velocity of GcKIF5A depends on the structural composition of microtubules.

**Figure 5.**
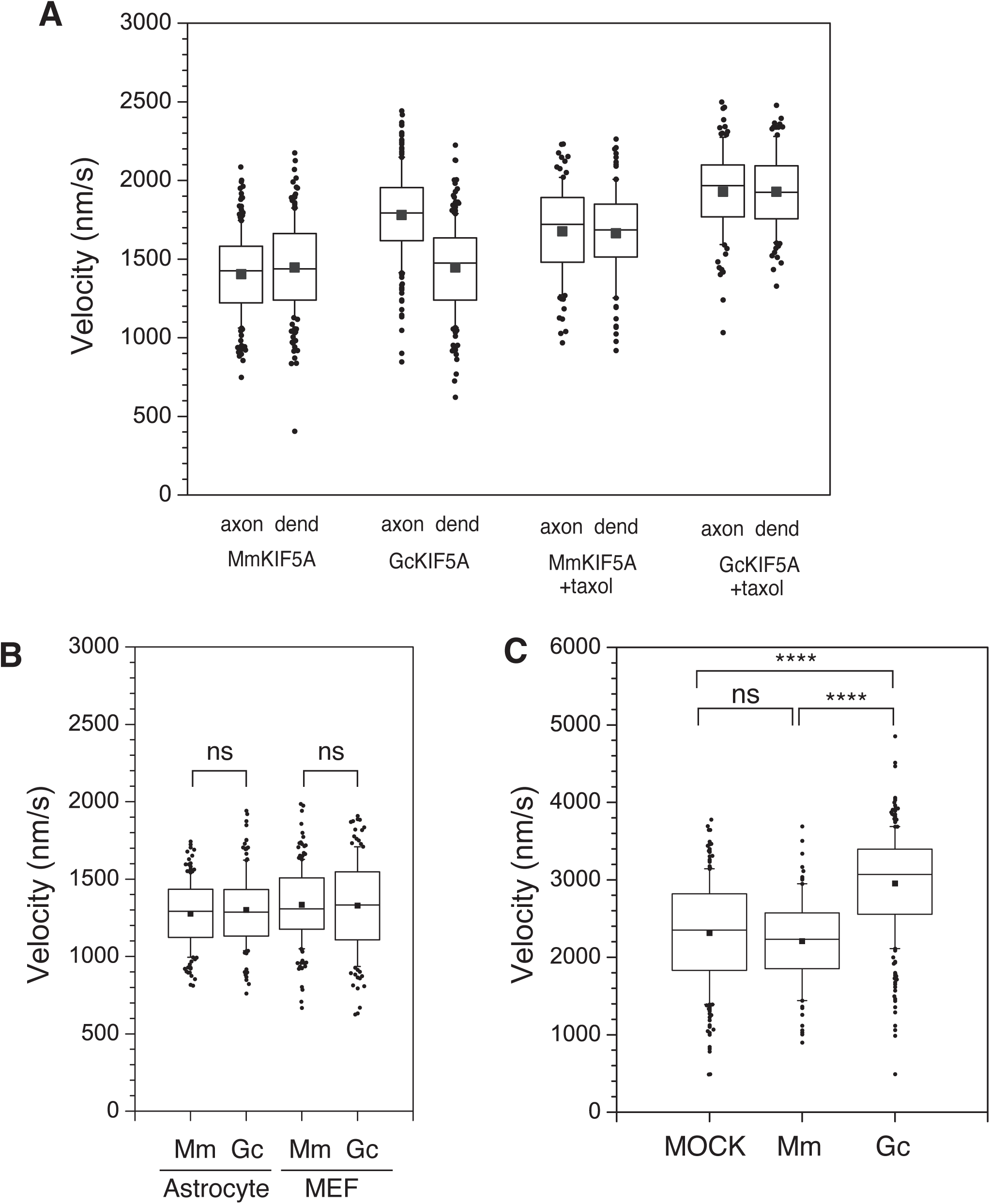
GcKIF5A Moves Faster in Axons but Not in Dendrites and Non-Neuronal Cells. (A) Comparison of K560-GFP motility in axons and dendrites of stage 3 primary mouse hippocampal neurons. K560 constructs of MmKIF5A or GcKIF5A were expressed in stage 3 neurons in the absence or presence of 1 μ*M* taxol. Box-and-whisker plots show the velocities of: MmKIF5A in axons (n=193 from 8 cells without taxol; n=100 from 4 cells with taxol), MmKIF5A in dendrites (n=185 from 8 cells without taxol; n=93 from 3 cells with taxol), GcKIF5A in axons (n=205 from 8 cells without taxol; n=100 from 4 cells with taxol), and GcKIF5A in dendrites (n=199 from 8 cells without taxol; n=100 from 3cells with taxol). (B) Velocity of GcKIF5A and MmKIF5A in non-neuronal cells. GcKIF5A and MmKIF5A were expressed in astrocyte or mouse embryonic fibroblasts (MEF). Velocities were measured in: MmKIF5A (n=153 from 4 cells in astrocyte; n=183 from 7 cells in MEF), and GcKIF5A (n=153 from 4 cells in astrocyte; n=144 from 7 cells in MEF). No significant differences were observed (Student’s t-test). (C) Transport of VSVG-RFP-positive vesicles by full-length KIF5A. Vesicle velocities by full-length KIF5A. Vesicles velocities were measured in cells with: No exogenous KIF5A (MOCK, n=232 from 23 cells), with expressed full-length MmKIF5A-EGFP (n=97 from 18 cells), and with expressed full-length GcKIF5A-EGFP (n=272 from 24 cells). Data are shown as box-whisker plots. ns, not significant; **** P < 0.0001 (Tukey’s test).

To further assess axonal transport in a physiologically relevant context, we monitored the anterograde movement of VSVG-containing vesicles, a well-established cargo for kinesin-1-mediated axonal transport. In cells transfected with an empty plasmid (MOCK) or a plasmid for full-length EGFP-MmKIF5A, VSVG vesicles moved at 2314 ± 44 nm/s and 2209 ± 58 nm/s, respectively. Expression of full-length GcKIF5A increased vesicle velocity to 2951 ± 40 nm/s— a 33% increase relative to MmKIF5A (**Figure 5C)**. As expected for a processive motor with high duty ratio and low velocity sensitivity at loads below ∼3 pN, vesicle velocity was independent of KIF5A expression level, likely due to load sharing among multiple motors (**Figure S10**).

### GcKIF5A Minimizes Drag in Mixed-Motor Transport and Enables Efficient Long-Distance Cargo Movement

Although GcKIF5A exhibits higher velocity than MmKIF5A, its run length is less than half (Fig. 2). If a single motor molecule were solely responsible for long-distance transport, this shorter run length would indeed be paradoxical for a motor presumed to support axonal transport over several meters. However, in actual cellular environments, multiple motor molecules might typically work in coordination to transport cargo over long distances^39,40^. In such multi-motor contexts, having a longer run length may not necessarily be advantageous. To explore this hypothesis, we investigated how GcKIF5A functions when working with other motors and in teams.

First, we asked whether the transport velocity is affected by motor cooperation when multiple motors simultaneously drive motility. Using microtubule-gliding assays with varying motor densities as a proxy for different levels of motor cooperation, we found that both GcKIF5A and PbKIF5A consistently moved faster than MmKIF5A across all densities, with no evidence of cooperative acceleration or deceleration as motor density increased (**Figure S11**). This result aligns with our cellular data, where VSVG vesicle velocity was independent of KIF5A expression level (**Figure S10**).

Next, we examined how GcKIF5A behaves when operating alongside faster motors. While force opposing the motor’s direction (hindering load) can slow movement, force in the direction of motion (assisting load) does not accelerate KIF5A^41,42^, as previously shown for KIF5 and KIF1A mixtures^43^. In such scenarios, even a minority of slower motors can reduce the overall velocity of transport. Indeed, just 20% KIF5 in a KIF1A motor mixture was sufficient to shift the microtubule-gliding velocity toward that of KIF1A^43^.

Because GcKIF5A has a higher microtubule detachment rate under load (3.9 s^−1^) than MmKIF5A (1.3 s^−1^), we hypothesized that it might exert less drag in mixed-motor conditions. To test this, we performed microtubule-gliding assays using defined mixtures of KIF5A and the faster KIF1A (**Figure 6A)**. Both GcKIF5A and MmKIF5A reduced the velocity of KIF1A-driven gliding in a dose-dependent manner, but the effect was milder for GcKIF5A at 10-20% of total motor input, indicating that GcKIF5A imparts less resistance than MmKIF5A when multiple motors simultaneously drive the same microtubule.

**Figure 6.**
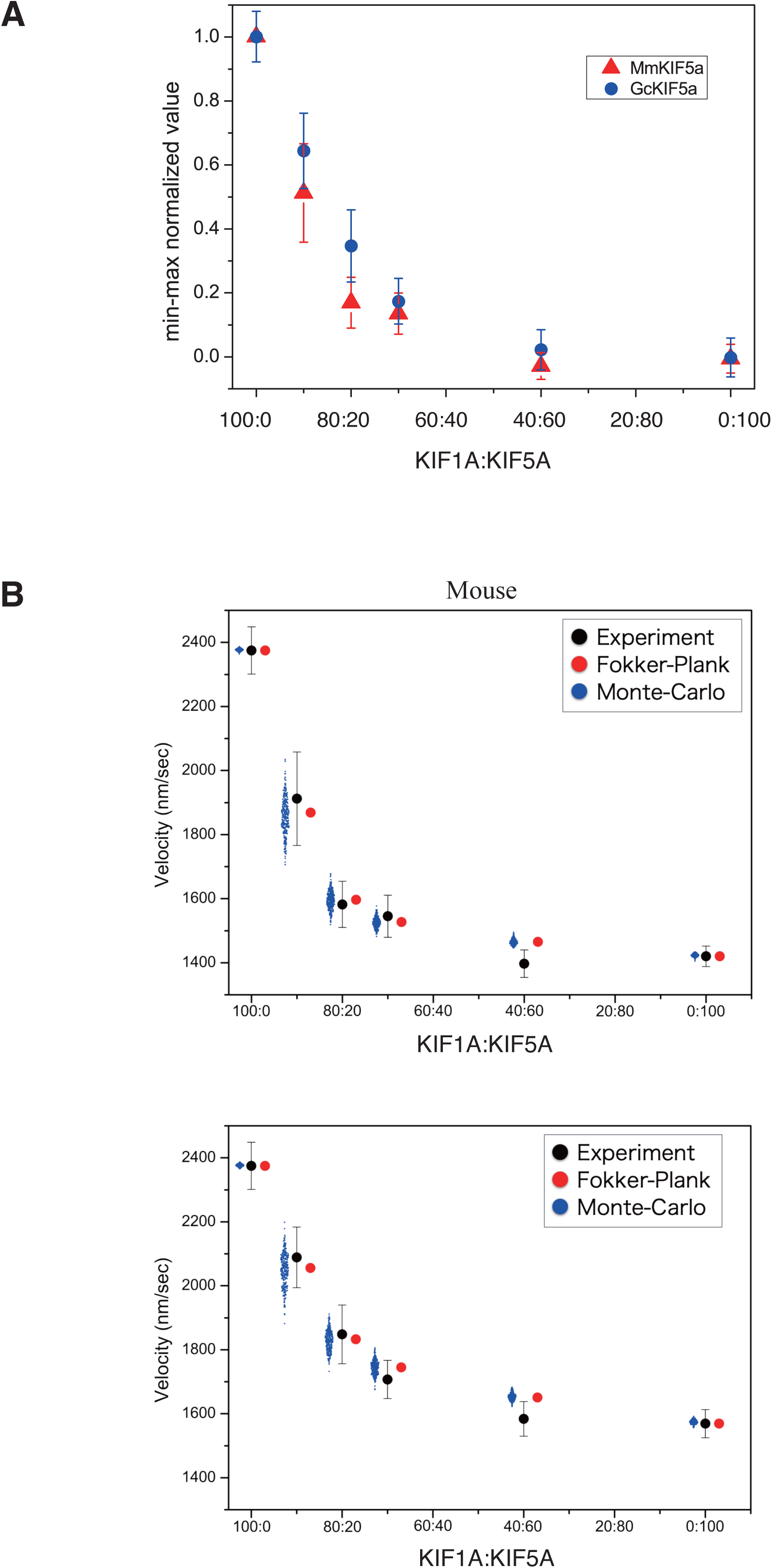
GcKIF5A Exhibits Reduced Drag in Mixed-Motor Transport, Supporting Faster Long-Distance Cargo Transport. (A) Microtubule-gliding assays using mixtures of KIF5A and the faster motor KIF1A at varying proportions. Gliding velocity was measured as a function of KIF5A content to assess its impact on overall transport speed. GcKIF5A caused less reduction in velocity than MmKIF5A, indicating reduced drag in mixed-motor conditions. (B) Computational simulations using both Fokker-Plank and Monte-Carlo approaches reproduce experimental data and reveal that differences in unloaded detachment rates account for the reduced drag effect of GcKIF5A. Parameters were based on previously published models.

To quantify these observations, we implemented a computational model of gliding assays based on parameters from Arpag et al.^43^. Simulations using Monte Carlo and Fokker-Planck approaches (**Figure 6B)** modeled the force-dependent detachment rate as 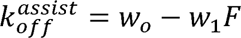, where w_0_ is the unloaded detachment rate and w_1_ is the slope of the force-detachment curve. The fitted *w*_1_ values were similar for both motors (0.9 ± 0.29 pN^−1^s^−1^ MmKIF5A and 0.76 ± 0.31 pN^−1^s^−1^ for GcKIF5A), suggesting that the reduced drag effect of GcKIF5A is primarily due to its elevated detachment rate in the absence of load, rather than altered load sensitivity.

Together, these results reveal that despite its shorter run length, GcKIF5A is well-suited for long-distance transport by rapidly moving cargo and minimizing drag when acting alongside other motors, such as KIF1A. This feature may be particularly advantageous in the context of large axonal geometries, where multiple motors are likely to coordinate cargo movement over exceptionally extended distances.

## DISCUSSION

In this study, we show that the kinesin motor KIF5A from giraffes—an animal with axons exceeding four meters in length—moves ∼25% faster than mouse KIF5A (MmKIF5A) on purified brain microtubules *in vitro* and within axons of mouse hippocampal neurons. While this 25% increase may appear modest relative to the dramatic difference in axonal length between giraffes and mice (nearly 100-fold), such velocity enhancements would confer substantial benefits to transport efficiency over extended distances. Notably, human KIF5A moves at a velocity comparable to mouse KIF5A, despite humans having axons over ten times longer, suggesting that motor velocity would become rate-limiting only in species with axons exceeding one meter in length. This threshold is supported by clinical observations in hereditary spastic paraplegia (HSP), where KIF5A mutations causing ∼25% velocity reductions^15^ primarily affect the longest motor neurons, with symptoms typically manifesting first in distal leg muscles innervated by meter-long axons. These observations collectively support the idea that modest velocity increases could become physiologically significant specifically in animals with exceptionally elongated axons, such as giraffes. Additionally, our finding that GcKIF5A moves preferentially faster on axonal versus dendritic microtubules indicates an additional layer of adaptation, potentially involving selective engagement with specific post-translationally modified tubulin subsets that could predominate in long-projection axons.

The identical triple amino acid substitution in giraffe and python KIF5A suggests a case of convergent evolution, potentially driven by the shared requirement for long-distance axonal transport. Among the 151 species analyzed, only these two—which possess some of the longest axons—harbor all three substitutions simultaneously. Notably, while the complete set of mutations is unique to giraffes and pythons, we observed partial substitutions (one or two of these three residues) in large animals such as cetartiodactyls with neurons featuring axons exceeding one meter in length. This pattern of partial adaptation provides evolutionary corroboration for our hypothesis that axons longer than one meter benefit from enhanced KIF5A velocity. The correlation between KIF5A adaptation and axonal length, rather than body size alone, is further supported by the species distribution of these substitutions throughout our phylogenetic analysis.

The physiological role of faster KIF5A in large animals is particularly intriguing when considering specific cellular contexts. While steady-state cargo delivery might not be critically dependent on motor velocity, our data suggest that faster KIF5A would be advantageous during neuronal development, when rapid material transport could support the extensive growth of exceptionally long axons. This developmental context hypothesis is supported by our comparative analysis of KIF5 isoforms: unlike GcKIF5C, which did not exhibit faster motility than its mouse counterpart, GcKIF5A appears to have specifically evolved increased velocity. This functional specialization aligns with physiological roles of these isoforms: KIF5A would be essential for axon formation and neuron survival, whereas KIF5C would be required later for motor neuron maintenance^9,11^. The temporal expression patterns further support this distinction— the expression of KIF5C increases after the first postnatal week. Consequently, KIF5C knockout mice are viable at birth, but experience postnatal motor neuron loss^9^. In contrast, KIF5A knockout mice die shortly after birth with impaired axon outgrowth^11^. Collectively, these observations suggest that the increased velocity of GcKIF5A likely represents an evolutionary adaptation to meet the demanding transport requirements during development of neurons with extremely long projections.

Mechanistically, our data demonstrate that the triple substitution in GcKIF5A is both necessary and sufficient to increase motor velocity. Each mutation appears to contribute to the enhanced function through distinct mechanisms. Y309F had the strongest effect in unloaded conditions, whereas S155A emerged as the primary contributor under load. This functional divergence resembles previous findings in kinesins with weakened cover-neck bundle (CNB) interactions, which display reduced stall force, increased unloaded velocity, and heightened force sensitivity^44^.

Our structural studies illuminate the molecular basis for these contributions. Crystallographic analysis revealed that the nucleotide-binding pocket of GcKIF5A adopts a more open conformation favoring accelerated ADP release. The Y309F substitution, located at the pivot point of helix a6 near the neck-linker base (**Figure 4**), would likely alter CNB formation through tighter packing between P-loop, helix α1a, and helix α6 elements. Our cryo-EM structure showed that this arrangement positions helix α6 closer to the microtubule surface, enhancing its interaction with α-tubulin and promoting more efficient neck-linker docking and power stroke under no-load conditions. These structural changes explain why Y309F strongly enhances unloaded velocity yet shows negligible effects under force—a characteristic pattern consistent with altered CNB interactions^44^.

In contrast, S155A resides at the microtubule-binding interface within the β5a strand and does not affect helix a6 conformation (**Figure 4**), consistent with its minimal impact on unloaded velocity. However, this substitution reduces β5a-L8 density in our cryo-EM structure, indicating increased conformational flexibility that would likely modulate load-dependent stepping by altering microtubule interactions under tension. While this region does not directly participate in ATP hydrolysis, its orientation toward the microtubule would influence how GcKIF5A responds to mechanical forces, especially in the crowded environment of the axon.

The third substitution, R114Q, positioned above the nucleotide-binding pocket, likely works synergistically with the other mutations to fine-tune the ADP release process that limits the overall catalytic cycle rate. Together, these structural adaptations collectively optimize GcKIF5A for its specialized role in long-distance axonal transport in large vertebrates.

Consistent with these structural insights, GcKIF5A maintained elevated velocity under both unloaded and loaded conditions. Under opposing forces up to ∼4 pN, GcKIF5A and the mouse triple mutant MmKIF5A retained velocities approximately 20% higher than wild-type mouse KIF5A (**Figure 3E)**. This robustness under load is essential for long-distance cargo transport in extremely long axons, meeting the unique demands of large animals like giraffes and pythons. Moreover, we show that GcKIF5A exerts less drag in mixed-motor conditions, likely due to its higher unloaded detachment rate. In cargo transport scenarios involving both fast (e.g., KIF1A) and slower motors, this property allows GcKIF5A to disengage more readily, avoiding the velocity-limiting effect observed with MmKIF5A. Together, these features enable GcKIF5A to support rapid, processive transport in the unique cellular architecture of large vertebrates.

In summary, our findings reveal a molecular adaptation of KIF5A that supports the extreme transport demands of neurons in large animals. The enhanced velocity, load resistance, and cooperative motor behavior of GcKIF5A provide insight into the evolutionary diversification of kinesin motors and underscore the importance of fine-tuning motor function to meet organismal-scale challenges posed by exceptionally elongated axons.

## RESOURCE AVAILABILITY

### Lead contact

Further information and requests for resources and reagents should be directed to and will be fulfilled by the lead contact, Yasushi Okada.

### Materials availability

Plasmids will be available through Addgene and RIKEN Bioresource Center (DNA Bank). All other unique/stable reagents generated in this study are available from the lead contact with a completed materials transfer agreement (MTA).

### Data and code availability

The atomic coordinates of the GcKIF5A and GcKIF5A-microtubule complex have been deposited in the PDB under accession codes 9UMM, 9UMN, and 9UMU, respectively. Electron microscopy maps are available in the Electron Microscopy Data Bank with accession code EMD-64332. Quantitative cell biological data will be in the SSBD database at https://ssbd.qbic.riken.jp/ with ID ssbd-repos-000433. Any additional information required to reanalyze the data reported in this paper is available from the lead contact upon request.

## ACKNOWLEDGMENTS

We thank Dr. Shinsuke Niwa for helpful discussions. We also thank our lab members for their discussion and support, especially Ms. Xu Shang Dan, Yukiko Onishi, Mikako Hayashi, Sayuri Yamamoto, and Junko Asada for technical support, and Ms. Manaho Kakiuchi, Ryoko Araki, and Tomoko Furuya for secretarial assistance. This work was supported by Japan Society for the Promotion of Science (JSPS) KAKENHI grants (JP19H03394, JP19H05794, JP19H05795, JP22H02798 to Y.O. and T.K.; JP22H04926 to Y.O.; JP21H05254, JP23K24057 to R.N.; 22K06809 and 25H02379 to T. I.); Japan Science and Technology Agency (JST) CREST grants (JPMJCR1852, JPMJCR20E2 and JPMJCR24T2 to Y.O.); JST Moonshot Research and Development Program grant (JPMJMS2025-14 to Y.O.; JPMJMS2024-7 to R.N.); JST FOREST (JPMJFR214K to T. I.); Japan Agency for Medical Research and Development (AMED) CREST grant (JP23gm1710001s0502 to Y.O. and T.K.; JP21gm161003 to T. I.); National Institute of Health (R01GM147332 to L.R. and A.G.).

## DECLARATION OF GENERATIVE AI AND AI-ASSISTED TECHNOLOGIES

During the preparation of this work, the authors used Claude 3.7 Sonnet to improve the readability and language of the manuscript. These tools were utilized specifically for checking terminology consistency, unifying formatting, and enhancing overall language fluency. After using these services, the authors reviewed and edited the content as needed and take full responsibility for the content of the publication.

## SUPPLEMENTAL INFORMATION

Materials and Methods

Supplemental Figures S1-S10

Supplemental Tables S1, S2

## REFERENCES

1. Guedes-Dias, P., and Holzbaur, E.L.F. (2019). Axonal transport: Driving synaptic function. Science 366.

2. De Vos, K., Grierson, A., Ackerley, S., and Miller, C. (2008). Role of Axonal Transport in Neurodegenerative Diseases. Annu. Rev. Neurosci. 31, 151–173.

3. De Vos, K.J., and Hafezparast, M. (2017). Neurobiology of axonal transport defects in motor neuron diseases: Opportunities for translational research? Neurobiol. Dis. 105, 283–299.

4. Coleman, M.P., and Hoke, A. (2020). Programmed axon degeneration: from mouse to mechanism to medicine. Nat. Rev. Neurosci. 21, 183–196.

5. Millecamps, S., and Julien, J.P. (2013). Axonal transport deficits and neurodegenerative diseases. Nat. Rev. Neurosci. 14, 161–176.

6. Sleigh, J.N., Rossor, A.M., Fellows, A.D., Tosolini, A.P., and Schiavo, G. (2019). Axonal transport and neurological disease. Nat. Rev. Neurol. 15, 691–703.

7. Hirokawa, N., Noda, Y., and Okada, Y. (1998). Kinesin and dynein superfamily proteins in organelle transport and cell division. Curr. Opin. Cell Biol. 10, 60–73.

8. Hirokawa, N., Niwa, S., and Tanaka, Y. (2010). Molecular Motors in Neurons: Transport Mechanisms and Roles in Brain Function, Development, and Disease. Neuron 68, 610–638.

9. Kanai, Y., Okada, Y., Tanaka, Y., Harada, A., Terada, S., and Hirokawa, N. (2000). KIF5C, a novel neuronal kinesin enriched in motor neurons. J. Neurosci. 20, 6374–6384.

10. Miki, H., Setou, M., Kaneshiro, K., and Hirokawa, N. (2001). All kinesin superfamily protein, KIF, genes in mouse and human. Proc. Natl. Acad. Sci. USA 98, 7004–7011.

11. Karle, K.N., Mockel, D., Reid, E., and Schols, L. (2012). Axonal transport deficit in a KIF5A(-/-) mouse model. Neurogenetics 13, 169–179.

12. Tanaka, Y., Kanai, Y., Okada, Y., Nonaka, S., Takeda, S., Harada, A., and Hirokawa, N. (1998). Targeted disruption of mouse conventional kinesin heavy chain, kif5B, results in abnormal perinuclear clustering of mitochondria. Cell 93, 1147–1158.

13. Xia, C.H., Roberts, E.A., Her, L.S., Liu, X., Williams, D.S., Cleveland, D.W., and Goldstein, L.S. (2003). Abnormal neurofilament transport caused by targeted disruption of neuronal kinesin heavy chain KIF5A. J. Cell Biol. 161, 55–66.

14. Reid, E., Kloos, M., Ashley-Koch, A., Hughes, L., Bevan, S., Svenson, I.K., Graham, F.L., Gaskell, P.C., Dearlove, A., Pericak-Vance, M.A., et al. (2002). A kinesin heavy chain (KIF5A) mutation in hereditary spastic paraplegia (SPG10). Am. J. Hum. Genet. 71, 1189–1194.

15. Dutta, M., Diehl, M.R., Onuchic, J.N., and Jana, B. (2018). Structural consequences of hereditary spastic paraplegia disease-related mutations in kinesin. Proc. Natl. Acad. Sci. USA 115, E10822–E10829.

16. Ebbing, B., Mann, K., Starosta, A., Jaud, J., Schöls, L., Schüle, R., and Woehlke, G. (2008). Effect of spastic paraplegia mutations in KIF5A kinesin on transport activity. Hum. Mol. Genet. 17, 1245–1252.

17. Crimella, C., Baschirotto, C., Arnoldi, A., Tonelli, A., Tenderini, E., Airoldi, G., Martinuzzi, A., Trabacca, A., Losito, L., Scarlato, M., et al. (2012). Mutations in the motor and stalk domains of KIF5A in spastic paraplegia type 10 and in axonal Charcot-Marie-Tooth type 2. Clin. Genet. 82, 157–164.

18. Nam, D.E., Yoo, D.H., Choi, S.S., Choi, B.O., and Chung, K.W. (2018). Wide phenotypic spectrum in axonal Charcot-Marie-Tooth neuropathy type 2 patients with KIF5A mutations. Genes Genomics 40, 77–84.

19. Brenner, D., Yilmaz, R., Muller, K., Grehl, T., Petri, S., Meyer, T., Grosskreutz, J., Weydt, P., Ruf, W., Neuwirth, C., et al. (2018). Hot-spot KIF5A mutations cause familial ALS. Brain 141, 688–697.

20. Nicolas, A., Kenna, K.P., Renton, A.E., Ticozzi, N., Faghri, F., Chia, R., Dominov, J.A., Kenna, B.J., Nalls, M.A., Keagle, P., et al. (2018). Genome-wide Analyses Identify KIF5A as a Novel ALS Gene. Neuron 97, 1267–1288.

21. Fichera, M., Lo Giudice, M., Falco, M., Sturnio, M., Amata, S., Calabrese, O., Bigoni, S., Calzolari, E., and Neri, M. (2004). Evidence of kinesin heavy chain (KIF5A) involvement in pure hereditary spastic paraplegia. Neurology 63, 1108–1110.

22. Lo Giudice, M., Neri, M., Falco, M., Sturnio, M., Calzolari, E., Di Benedetto, D., and Fichera, M. (2006). A missense mutation in the coiled-coil domain of the KIF5A gene and late-onset hereditary spastic paraplegia. Arch. Neurol. 63, 284–287.

23. Blair, M.A., Ma, S., and Hedera, P. (2006). Mutation in KIF5A can also cause adult-onset hereditary spastic paraplegia. Neurogenetics 7, 47–50.

24. Wilkinson, D.M., and Ruxton, G.D. (2012). Understanding selection for long necks in different taxa. Biol. Rev. Camb. Philos. Soc. 87, 616–630.

25. Agaba, M., Ishengoma, E., Miller, W.C., McGrath, B.C., Hudson, C.N., Bedoya Reina, O.C., Ratan, A., Burhans, R., Chikhi, R., Medvedev, P., et al. (2016). Giraffe genome sequence reveals clues to its unique morphology and physiology. Nat. Commun. 7, 11519.

26. Seymour, R.S., and Johansen, K. (1987). Blood flow uphill and downhill: does a siphon facilitate circulation above the heart? Comp. Biochem. Physiol. A Comp. Physiol. 88, 167–170.

27. Ferro, L.S., Fang, Q., Eshun-Wilson, L., Fernandes, J., Jack, A., Farrell, D.P., Golcuk, M., Huijben, T., Costa, K., Gur, M., et al. (2022). Structural and functional insight into regulation of kinesin-1 by microtubule-associated protein MAP7. Science 375, 326–331.

28. Kuimova, M.K., Yahioglu, G., Levitt, J.A., and Suhling, K. (2008). Molecular rotor measures viscosity of live cells via fluorescence lifetime imaging. J. Am. Chem. Soc. 130, 6672–6673.

29. Luby-Phelps, K., Mujumdar, S., Mujumdar, R.B., Ernst, L.A., Galbraith, W., and Waggoner, A.S. (1993). A novel fluorescence ratiometric method confirms the low solvent viscosity of the cytoplasm. Biophys. J. 65, 236–242.

30. Suhling, K., Siegel, J., Lanigan, P.M., Leveque-Fort, S., Webb, S.E., Phillips, D., Davis, D.M., and French, P.M. (2004). Time-resolved fluorescence anisotropy imaging applied to live cells. Opt. Lett. 29, 584–586.

31. Fushimi, K., and Verkman, A.S. (1991). Low viscosity in the aqueous domain of cell cytoplasm measured by picosecond polarization microfluorimetry. J. Cell Biol. 112, 719–725.

32. Cao, L., Cantos-Fernandes, S., and Gigant, B. (2017). The structural switch of nucleotide-free kinesin. Sci. Rep. 7, 42558.

33. Morikawa, M., Yajima, H., Nitta, R., Inoue, S., Ogura, T., Sato, C., and Hirokawa, N. (2015). X-ray and Cryo-EM structures reveal mutual conformational changes of Kinesin and GTP-state microtubules upon binding. EMBO J. 34, 1270–1286.

34. Ma, Y.Z., and Taylor, E.W. (1997). Kinetic mechanism of a monomeric kinesin construct. J. Biol. Chem. 272, 717–723.

35. Nakata, T., Niwa, S., Okada, Y., Perez, F., and Hirokawa, N. (2011). Preferential binding of a kinesin-1 motor to GTP-tubulin-rich microtubules underlies polarized vesicle transport. J. Cell Biol. 194, 245–255.

36. Janke, C. (2014). The tubulin code: molecular components, readout mechanisms, and functions. J. Cell Biol. 206, 461–472.

37. Shima, T., Morikawa, M., Kaneshiro, J., Kambara, T., Kamimura, S., Yagi, T., Iwamoto, H., Uemura, S., Shigematsu, H., Shirouzu, M., et al. (2018). Kinesin-binding-triggered conformation switching of microtubules contributes to polarized transport. J. Cell Biol. 217, 4164–4183.

38. Janke, C., and Magiera, M.M. (2020). The tubulin code and its role in controlling microtubule properties and functions. Nat. Rev. Mol. Cell Biol. 21, 307–326.

39. Hayashi, K., Hasegawa, S., Sagawa, T., Tasaki, S., and Niwa, S. (2018). Non-invasive force measurement reveals the number of active kinesins on a synaptic vesicle precursor in axonal transport regulated by ARL-8. Phys. Chem. Chem. Phys. 20, 3403–3410.

40. Hayashi, K., Matsumoto, S., Naoi, T., and Idobata, Y. (2021). Intracellular force comparison of pathogenic KIF1A, KIF5, and dynein by fluctuation analysis. Preprint at bioRxiv, 2021.09.12.459977.

41. Andreasson, J.O., Milic, B., Chen, G.Y., Guydosh, N.R., Hancock, W.O., and Block, S.M. (2015). Examining kinesin processivity within a general gating framework. Elife 4, e07403.

42. Block, S.M., Asbury, C.L., Shaevitz, J.W., and Lang, M.J. (2003). Probing the kinesin reaction cycle with a 2D optical force clamp. Proc. Natl. Acad. Sci. USA 100, 2351–2356.

43. Arpag, G., Shastry, S., Hancock, W.O., and Tuzel, E. (2014). Transport by populations of fast and slow kinesins uncovers novel family-dependent motor characteristics important for in vivo function. Biophys. J. 107, 1896–1904.

44. Khalil, A.S., Appleyard, D.C., Labno, A.K., Georges, A., Karplus, M., Belcher, A.M., Hwang, W., and Lang, M.J. (2008). Kinesin’s cover-neck bundle folds forward to generate force. Proc. Natl. Acad. Sci. USA 105, 19247–19252.

45. Altschul, S.F., Madden, T.L., Schaffer, A.A., Zhang, J., Zhang, Z., Miller, W., and Lipman, D.J. (1997). Gapped BLAST and PSI-BLAST: a new generation of protein database search programs. Nucleic Acids Res. 25, 3389–3402.

46. Castoldi, M., and Popov, A.V. (2003). Purification of brain tubulin through two cycles of polymerization-depolymerization in a high-molarity buffer. Protein Expr. Purif. 32, 83–88.

47. Friel, C.T., and Howard, J. (2011). The kinesin-13 MCAK has an unconventional ATPase cycle adapted for microtubule depolymerization. EMBO J. 30, 3928–3939.

48. Furuta, K., and Toyoshima, Y.Y. (2008). Minus-end-directed motor Ncd exhibits processive movement that is enhanced by microtubule bundling in vitro. Curr. Biol. 18, 152–157.

49. Jiang, M., and Chen, G. (2006). High Ca2+-phosphate transfection efficiency in low-density neuronal cultures. Nat. Protoc. 1, 695–700.

50. Nakata, T., and Hirokawa, N. (1995). Point mutation of adenosine triphosphate-binding motif generated rigor kinesin that selectively blocks anterograde lysosome membrane transport. J. Cell Biol. 131, 1039–1053.

51. Kabsch, W. (2010). Xds. Acta Crystallogr. D Biol. Crystallogr. 66, 125–132.

52. Emsley, P., Lohkamp, B., Scott, W.G., and Cowtan, K. (2010). Features and development of Coot. Acta Crystallogr. D Biol. Crystallogr. 66, 486–501.

53. Liebschner, D., Afonine, P.V., Baker, M.L., Bunkóczi, G., Chen, V.B., Croll, T.I., Hintze, B., Hung, L.W., Jain, S., McCoy, A.J., et al. (2019). Macromolecular structure determination using X-rays, neutrons and electrons: recent developments in Phenix. Acta Crystallogr. D Struct. Biol. 75, 861–877.

54. Pettersen, E.F., Goddard, T.D., Huang, C.C., Couch, G.S., Greenblatt, D.M., Meng, E.C., and Ferrin, T.E. (2004). UCSF Chimera--a visualization system for exploratory research and analysis. J. Comput. Chem. 25, 1605–1612.

55. Mastronarde, D.N. (2005). Automated electron microscope tomography using robust prediction of specimen movements. J. Struct. Biol. 152, 36–51.

56. Zivanov, J., Nakane, T., Forsberg, B.O., Kimanius, D., Hagen, W.J., Lindahl, E., and Scheres, S.H. (2018). New tools for automated high-resolution cryo-EM structure determination in RELION-3. Elife 7, e42166.

57. Zheng, S.Q., Palovcak, E., Armache, J.P., Verba, K.A., Cheng, Y., and Agard, D.A. (2017). MotionCor2: anisotropic correction of beam-induced motion for improved cryo-electron microscopy. Nat. Methods 14, 331–332.

58. Rohou, A., and Grigorieff, N. (2015). CTFFIND4: Fast and accurate defocus estimation from electron micrographs. J. Struct. Biol. 192, 216–221.

59. Lacey, S.E., He, S., Scheres, S.H., and Carter, A.P. (2019). Cryo-EM of dynein microtubule-binding domains shows how an axonemal dynein distorts the microtubule. Elife 8, e47145.

60. Cook, A.D., Manka, S.W., Wang, S., Moores, C.A., and Atherton, J. (2020). A microtubule RELION-based pipeline for cryo-EM image processing. J. Struct. Biol. 209, 107402.

61. Sui, H., and Downing, K.H. (2010). Structural basis of interprotofilament interaction and lateral deformation of microtubules. Structure 18, 1022–1031.

